# Metabolic engineering of a tyrosine-specific phenylpropanoid pathway in plants

**DOI:** 10.64898/2025.12.16.694581

**Authors:** Caroline Van Beirs, Max Bentelspacher, Congwei Xie, Celien Van de Velde, Sandrien Desmet, Robin De Wulf, Wout Boerjan, Jaime Barros-Rios, Bartel Vanholme

## Abstract

While all plants use L-phenylalanine for phenylpropanoid biosynthesis, grasses can also initiate the pathway from L-tyrosine. Curiously, no plant has evolved an exclusive tyrosine-derived route. We generate plants with phenylpropanoid biosynthesis initiated from phenylalanine, tyrosine, or both by expressing a *Brachypodium* phenylalanine/tyrosine ammonia-lyase (*PTAL)* in Arabidopsis WT and *c4h* mutants. Engineering a bifunctional phenylpropanoid pathway in WT plants did not negatively impact growth, while introducing a tyrosine-specific pathway in the *c4h* mutant could overcome the seedling-lethal phenotype. Interestingly, restored *c4h* mutants relying solely on the tyrosine route displayed developmental defects linked to the strong overaccumulation of the auxin transport inhibitor *cis*-cinnamic acid. Our findings suggest that the requirement of this widely overlooked plant metabolite could be the crucial factor for the evolutionary retention of the canonical phenylpropanoid biosynthesis route via L-phenylalanine in plants.

## INTRODUCTION

The phenylpropanoid pathway is essential for plant metabolism, as it leads to a broad range of aromatic metabolites (Vogt 2010). The end-products of the pathway seve as building blocks for lignin, mainly found in the secondary cell walls of plants and one of the most abundant natural polymers in the biosphere. It provides structural support and hydrophobicity essential for water transport and plant growth, making it an essential evolutionary invention for the diversification and success of land plants (Vogt 2010; Boerjan, Ralph, and Baucher 2003). In *Arabidopsis thaliana*, the first step of the phenylpropanoid pathway is performed by PHENYLALANINE AMMONIA LYASE (PAL), which catalyses the deamination of L-phenylalanine (Phe) to form *trans*-cinnamic acid (*trans*-CA, **Fig. 1A**). Afterwards, *trans-*CA is hydroxylated by CINNAMATE-4-HYDROXYLASE (C4H) to form *p*-coumaric acid, which is subsequently activated by 4-COUMARATE-CoA LIGASE (4CL) to form *p*-coumaroyl-CoA. Through a series of enzymatic reactions, *p*-coumaroyl-CoA is converted to the primary monolignols *p*-coumaryl alcohol, coniferyl alcohol, and sinapyl alcohol. These monolignols polymerize in the cell wall via radical coupling, producing the *p*-hydroxyphenyl (H), guaiacyl (G), and syringyl (S) units in the lignin polymer (Boerjan, Ralph, and Baucher 2003).

**Figure 1:**
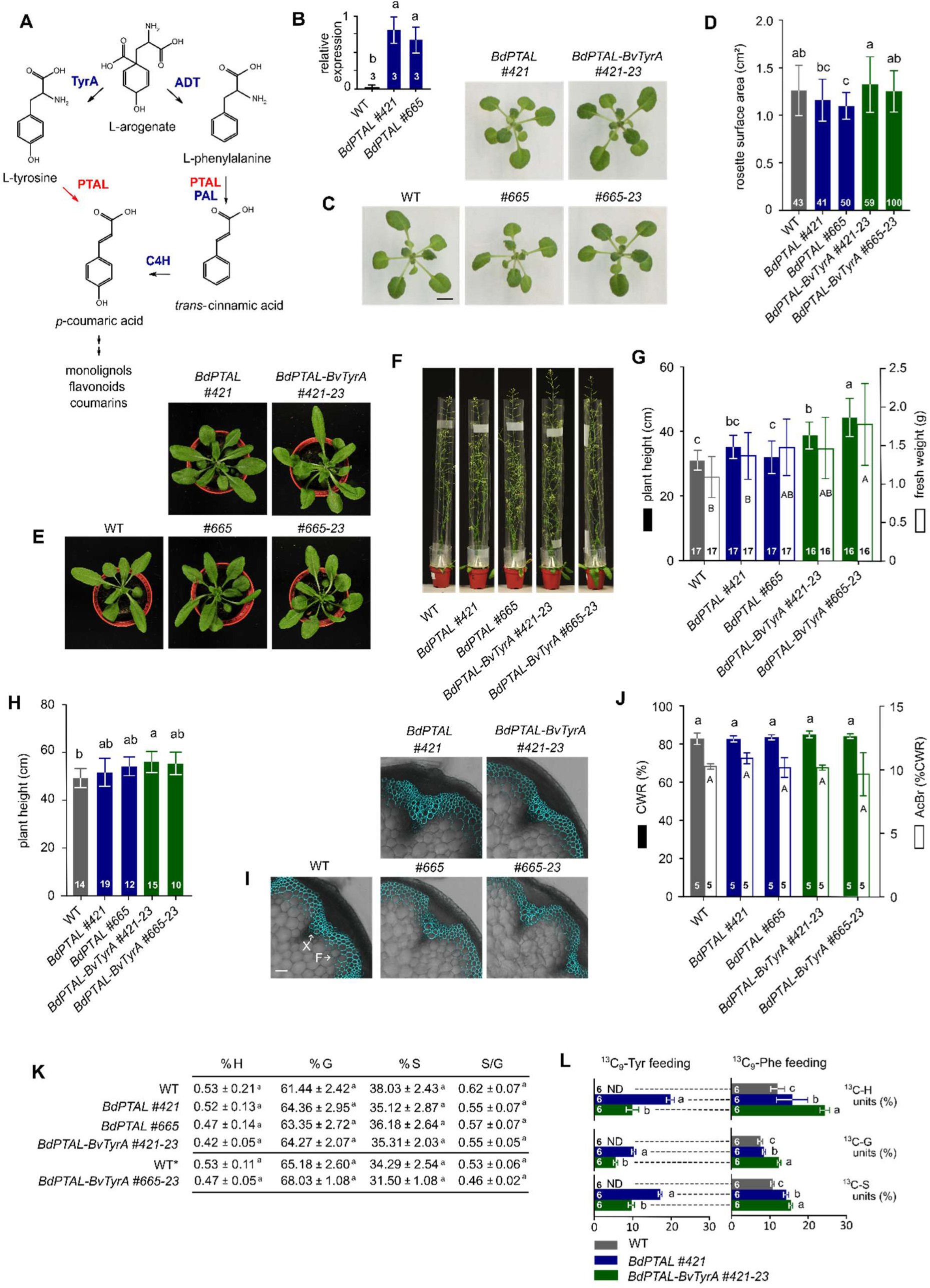
Introducing a dual phenylpropanoid pathway in Arabidopsis. (**A**) Intersection of the aromatic amino acid biosynthetic pathway and the early phenylpropanoid pathway. Enzymes are indicated in blue, with the exception of PTAL, which is indicated in red. (**B**) Relative expression of *BdPTAL* compared to housekeeping gene *UBC21* (At5g25760) in WT and the two selected *BdPTAL* overexpression lines assayed by qPCR (*n* =3). (**C**) Rosettes of *in vitro* grown WT and *BdPTAL* overexpression lines 21 DAS. Bar = 1 cm. (**D**) Projected rosette area of *in vitro* grown WT and *BdPTAL* overexpression lines 21 DAS. (**E**) Rosettes of 4-week-old soil-grown WT and *BdPTAL* overexpression lines. Pot diameter is 5.5 cm. (**F**) Inflorescence stems of 7-week-old WT and *BdPTAL* overexpression lines. Pot diameter is 5.5 cm. (**G**) Plant height and biomass of above-ground plant tissue (rosette + stem) of soil-grown WT and *BdPTAL* overexpression lines, 7 weeks after sowing. (**H**) Plant height of fully senesced WT and *BdPTAL* overexpression lines. (**I**) Lignin autofluorescence in the inflorescence stem of WT and *BdPTAL* overexpression lines, 7 weeks after sowing. Bar = 50 µm. F: interfascicular fibers, X: xylem. (**J**) CWR and lignin content of the lower part of the fully senesced stem of WT and *BdPTAL* overexpression lines. The CWR determined gravimetrically after sequential extraction and expressed as a percentage of dry weight. The lignin content was assayed by the AcBr method and expressed as a percentage of CWR. (**K**) Lignin composition of the CWR of WT and *BdPTAL* overexpression lines determined with thioacidolysis followed by GC-MC analysis. The relative proportions of the different lignin units were calculated based on the total thioacidolysis yield. S/G was calculated based on the absolute values for S and G. Values are given as average ± statistical deviation (*n*=5). Thioacidolysis for *BdPTAL-BvTyrA #665-23* was performed as an additional independent experiment, using *WT as an internal control. (**L**) Proportion of labeled precursors incorporated into each of the three main lignin units (H, G, and S) in 21-day-old *in vitro* grown plants grown on MS medium with isotopically labeled phenylalanine or tyrosine. The relative proportions of the different lignin units were calculated based on the total thioacidolysis yield (100% is the total area under the curve; n=6). Statistical significance was determined using an Anova test followed by a Tukey posthoc test. Numbers in the bars represent the number of repeats. Equal letters represent no statistical difference at the 0.05 significance level. Error bars represent standard deviation. AcBr: acetylbromide / ADT: AROGENATE DEHYDRATASE / BdPTAL: *Brachypodium distachyon* PHENYLALANINE TYROSINE AMMONIA LYASE / BvTyrA: *Brachypodium distachyon* AROGENATE DEHYDROGENASE / C4H: CINNAMATE-4-HYDROXYLASE / CWR: cell wall residue / DAS: days after stratification / ND: not detected / PAL: PHENYLALANINE AMMONIA LYASE / Phe: phenylalanine / PTAL: PHENYLALANINE TYROSINE AMMONIA LYASE / Tyr: tyrosine / TyrA: AROGENATE DEHYDROGENASE / WT: wild type

Beyond serving as precursors for lignin biosynthesis, several intermediates of the phenylpropanoid pathway are metabolized into compounds that participate in a broad range of biochemical and physiological processes. For example, *p*-coumarate acts as a key precursor for flavonoids, which function as antioxidants (Li et al. 2022); coumarins, which facilitate iron mobilization (Stringlis, De Jonge, and Pieterse 2019); and stilbenes, which serve as phytoalexins that enhance resistance to pathogenic attack (Al-Khayri et al. 2023). In addition, the precursor of *p*-coumarate, *trans*-CA, can convert into its *cis*-isomer which acts as a non-competitive inhibitor of auxin efflux carriers, thereby impacting auxin homeostasis (Steenackers et al. 2016; El Houari et al. 2021). These examples illustrate the versatility of phenylpropanoid metabolism and its central role in coordinating plant responses to developmental and environmental changes.

Besides PAL, grasses in the Poaceae family possess bifunctional PHENYLALANINE TYROSINE AMMONIA LYASES (PTALs) that can deaminate both L-tyrosine (Tyr) and Phe (Barros et al. 2016; Feduraev et al. 2020). Although Phe and Tyr both originate from L-arogenate, they are synthesized by two distinct enzymes: AROGENATE DEHYDROGENASE (TyrA/ADH) produces Tyr, while AROGENATE DEHYDRATASE (ADT) produces Phe (Schenck and Maeda 2018) (**Fig. 1A**). Metabolically, it is intriguing that all plants except grasses rely solely on Phe for phenylpropanoid biosynthesis, given that its synthesis involves the removal of the *para*-hydroxy group, which must later be reintroduced by C4H to form *p*-coumaric acid. In contrast, the Tyr route retains this hydroxyl group throughout, leading to a potentially more direct path to phenylpropanoid intermediates. Unlike bacteria, which can use monofunctional TYROSINE AMMONIA-LYASES (TALs) for producing certain chromophores and antibiotics (Berner et al. 2006; Kyndt et al. 2002), plants have never evolved an exclusive Tyr-specific pathway, suggesting metabolic constraints that make this route less favourable than the canonical Phe-derived phenylpropanoid pathway.

To explore the evolutionary significance of a Tyr-based phenylpropanoid pathway, we introduced a bifunctional *PTAL* from *Brachypodium distachyon* (*BdPTAL*) into *Arabidopsis thaliana*. Specifically, we transformed a heterozygous *C4H/c4h* mutant with a *BdPTAL* overexpression construct under the control of the Arabidopsis *C4H* promoter. Segregating progeny allowed the analysis of *BdPTAL* function in both WT and *c4h* mutant backgrounds. In a WT background, the expression of *BdPTAL* generated a dual-entry phenylpropanoid pathway via both Phe and Tyr. In contrast, in the *c4h* background, where the conversion of *trans*-CA to *p*-coumarate is blocked, all carbon skeletons channelled over the phenylpropanoid pathway originated from Tyr. Interestingly, whereas the *c4h* mutant is seedling lethal, the generated lines produced viable seeds under conditions where Tyr limitation was circumvented. In addition to the growth restoration, the plants showed auxin-related developmental defects including curled leaves, swollen branching sites and an accumulation of root hairs and lateral roots that were linked to the overaccumulation of *cis*-CA. Our findings suggest that the cellular demand for CA may impose a significant evolutionary constraint on the emergence of a dedicated monofunctional TAL pathway in plants.

## RESULTS

### Engineering a bifunctional PAL/TAL pathway in Arabidopsis

To investigate the consequences of engineering a grass-like dual PAL/TAL phenylpropanoid pathway in a dicot, we studied Arabidopsis plants overexpressing a bifunctional PTAL from the model grass *B. distachyon* (*BdPTAL*). The full-length *BdPTAL* (XP_003575396) coding sequence was cloned in an expression vector under the control of the Arabidopsis *C4H* promoter. After sequence confirmation, the construct was used to transform heterozygous *C4H/c4h* Arabidopsis plants using the floral dip method. At this point, progeny plants homozygous for *c4h* were excluded from further analysis, as we were first interested in studying plants with a dual phenylpropanoid pathway characteristic of monocot grass species. Plants expressing *BdPTAL* in either the WT or heterozygous *C4H*/*c4h* background were grouped for subsequent analysis, as WT and *C4H*/*c4h* are phenotypically indistinguishable (El Houari et al. 2021). Over three generations, two independent homozygous single *BdPTAL* insert lines with high *BdPTAL* expression were selected in a *C4H*/*c4h* heterozygous background (*BdPTAL* #421 and #665; **Fig. 1B**). These lines, along with WT control plants, were grown on 1/2× MS medium, and the projected rosette area was measured 21 days after stratification (DAS). WT plants had an average rosette surface area of 1.26 cm^2^, while *BdPTAL* #665 showed slightly smaller rosettes of 1.10 cm^2^, corresponding to a 12.9% decrease in rosette size compared to WT plants (**Fig. 1C, D**). Although *BdPTAL* #421 also showed a smaller average rosette size (1.16 cm^2^), the difference was not statistically significant from WT plants (*p* = 0.35).

The mild reduction in growth upon *BdPTAL* expression may be attributable to Tyr depletion. To mitigate this potential limitation, we pursued a genetic approach using the feedback-insensitive TyrA/ADH enzyme from *Beta vulgaris* (BvTyrA; **Fig. 1A**). Overexpression of the corresponding gene under the constitutive *35S* promoter in Arabidopsis has been shown to increase Tyr levels 60- to 300-fold (de Oliveira et al. 2019). Although these high Tyr levels have been linked to growth defects, *BvTyrA* line #23 showed no obvious phenotypes in our growth conditions (Supplementary Figure 1A-C). The *BvTyrA* #23 overexpression line was crossed with the two *C4H/c4h BdPTAL* lines, and progeny that were heterozygous for *C4H/c4h* and homozygous for *BdPTAL* and *BvTyrA* were selected and named *BdPTAL-BvTyrA* #421-23 and #665-23. When grown on 1/2× MS media, the *BdPTAL-BvTyrA* lines showed no reduction in rosette surface area compared to WT plants (**Fig. 1C, D**). Notably, both lines produced rosettes ∼12% larger than their corresponding *BdPTAL* overexpression controls. Plant growth was further assessed on soil, where no obvious visual differences in growth or development were observed between WT, *BdPTAL* or the *BdPTAL-BvTyrA* lines after four weeks of growth (**Fig. 1E)**. After seven weeks, both the *BdPTAL-BvTyrA* lines were slightly taller compared to WT (**Fig. 1F, G**), with *BdPTAL-BvTyrA* #665-23 showing the largest increase in plant height (30.7%). This line also showed a 62.6% increase in fresh biomass weight compared to WT plants. The growth difference diminished over the course of the growth season, and by senescence only the *BdPTAL-BvTyrA #421-23* line remained ∼12% taller than WT plants (**Fig. 1H)**. Together, these results indicate that engineering a dual PAL/TAL phenylpropanoid pathway in Arabidopsis is well tolerated and can even enhance growth relative to WT.

To evaluate the effect of *BdPTAL* and *BdPTAL-BvTyrA* expression on lignin deposition, cross-sections of inflorescence stems were analysed for lignin autofluorescence. WT controls showed round, open vessels, with lignin autofluorescence in the xylem and interfascicular fibres (**Fig. 1I**). *BdPTAL* and *BdPTAL-BvTyrA* overexpression lines showed similar autofluorescence patterns and vascular structure, indicating no major changes in lignin deposition and stem anatomy. Despite the absence of clear anatomical or histological effects, the overexpression of *BdPTAL* and *BdPTAL-BvTyrA* could have affected lignin content, which was evaluated through lignin analysis of the senesced primary inflorescence stem. Soluble metabolites were first removed by sequential chloroform/ethanol extraction, and the lignin content of the resulting cell wall residue (CWR) was quantified using the acetyl bromide (AcBr) method (Van Acker et al. 2013; Dence 1992). In line with the anatomical observation, no differences were found in the CWR and AcBr soluble lignin content between WT, *BdPTAL* and *BdPTAL-BvTyrA* overexpression lines (**Fig. 1J**). To assess lignin composition, the same stem tissues were analysed by thioacidolysis followed by gas chromatography coupled to mass spectrometry (GC-MS). This method quantifies the H, G, S lignin subunits that are linked by β-*O*-4 interunit bonds in the lignin polymer (Van Acker et al. 2013; Robinson and Mansfield 2009b). The amount of H, G, or S units released from the lignin of the *BdPTAL* or *BdPTAL-BvTyrA* lines were comparable to WT, and no significant changes were observed in the S/G ratio (**Fig. 1K**).

In the absence of a distinct lignin phenotype, we could not ascertain whether the Tyr-derived pathway contributed to the synthesis of monolignols. To verify the operation of the bifunctional PAL/TAL pathways and quantify the relative contributions of Phe and Tyr to lignin composition, we conducted stable isotopic tracing of phenylpropanoid biosynthesis using ^13^C_9_-labelled precursors on WT plants and both *BdPTAL #421* and *BdPTAL*-*BvTyrA #421-23* overexpression lines (**Fig. 1L**). Plants were grown on MS medium containing ^13^C_9_-Phe or ^13^C_9_-Tyr for three weeks and harvested for thioacidolysis followed by GC-MS analysis. Results showed that ^13^C_9_-Phe was incorporated into all lignin subunits (H, G, and S) in WT, as well as in *BdPTAL* and *BdPTAL-BvTyrA* lines. In contrast, ^13^C_9_-Tyr feeding resulted in no detectable label incorporation in WT plants, but substantial incorporation into all lignin subunits in both *BdPTAL* and *BdPTAL-BvTyrA* plants, indicating a functional TAL pathway in these lines. Considering the total incorporation of labeled units, 45 and 55% of the lignin originated from Phe or Tyr in line *BdPTAL* #421, values comparable to those reported in Brachypodium (**Supplementary Table 1**; Barros, 2016). Notably, the incorporation of ^13^C_9_-Tyr into the lignin subunits was significantly reduced in *BdPTAL-BvTyrA* lines compared to *BdPTAL* lines, suggesting that higher levels of endogenous Tyr pools in the *BdPTAL-BvTyrA* lines diluted the incorporation of ^13^C_9_-Tyr into lignin.

Taken together, these results show that parallel PAL and TAL pathways can be successfully engineered in dicot plants with minor effects on plant growth and lignin content or composition.

### Engineering a dedicated monofunctional TAL pathway

To test whether lignin biosynthesis can proceed exclusively from Tyr, we characterized the effects of *BdPTAL* and *BdPTAL-BvTyrA* constructs in the homozygous *c4h* knockout background. Arabidopsis *c4h* mutants display a seedling-lethal phenotype due to a complete block in phenylpropanoid biosynthesis (El Houari et al. 2021; Schilmiller et al. 2009). Restoration of the *c4h* phenotype by *BdPTAL* expression was considered indicative of a functional phenylpropanoid pathway exclusively derived from Tyr. The progeny of *C4H/c4h*, *C4H/c4h BdPTAL,* and *C4H/c4h BdPTAL-BvTyrA* plants were grown on horizontal plates alongside their respective controls, and the rosette surface areas were measured at 21 DAS (**Fig. 2A**). Homozygous *c4h* mutants were severely dwarfed and developed cotyledons but no true leaves, whereas the restored lines (*c4h BdPTAL* and *c4h BdPTAL-BvTyrA*) developed significantly larger and curled rosettes leaves, indicating that *BdPTAL*-overexpression can restore phenylpropanoid biosynthesis and rescue the growth phenotype of the *c4h* mutant. Despite the positive effect on plant growth, the restoration was only partial, with the rosette area of *c4h BdPTAL* #665 plants being still only 25.6% of that of WT plants (**Fig. 2B**). *BdPTAL* lines with lower transgene expression were not able to restore the *c4h* phenotype beyond the seedling stage (**Supplementary Figure 2A, B**). Adding the *BvTyrA* construct to both partly restored *c4h BdPTAL* lines further increased plant growth, though this increase was only significant for line #665. This data suggests that Tyr availability could be a limiting factor for the growth of plants with an exclusively Tyr-derived phenylpropanoid pathway. Supporting this observation, the rosette surface area increased when the *c4h BdPTAL* lines were grown on 1/2x MS medium supplemented with 50 µM Tyr (**Fig. 2C**). Although still additive, the effect of Tyr feeding was smaller in the *c4h BdPTAL* #421 line (24.1% increase) compared to the #665 line (68.0% increase). The difference between both lines could be due to the stronger complementation effect of the *c4h BdPTAL* #421 line, having already a project rosette area of 64.4% of the WT in the absence of exogenous supplied tyrosine, whereas this was only 18.1% for *c4h BdPTAL* #665.

**Figure 2:**
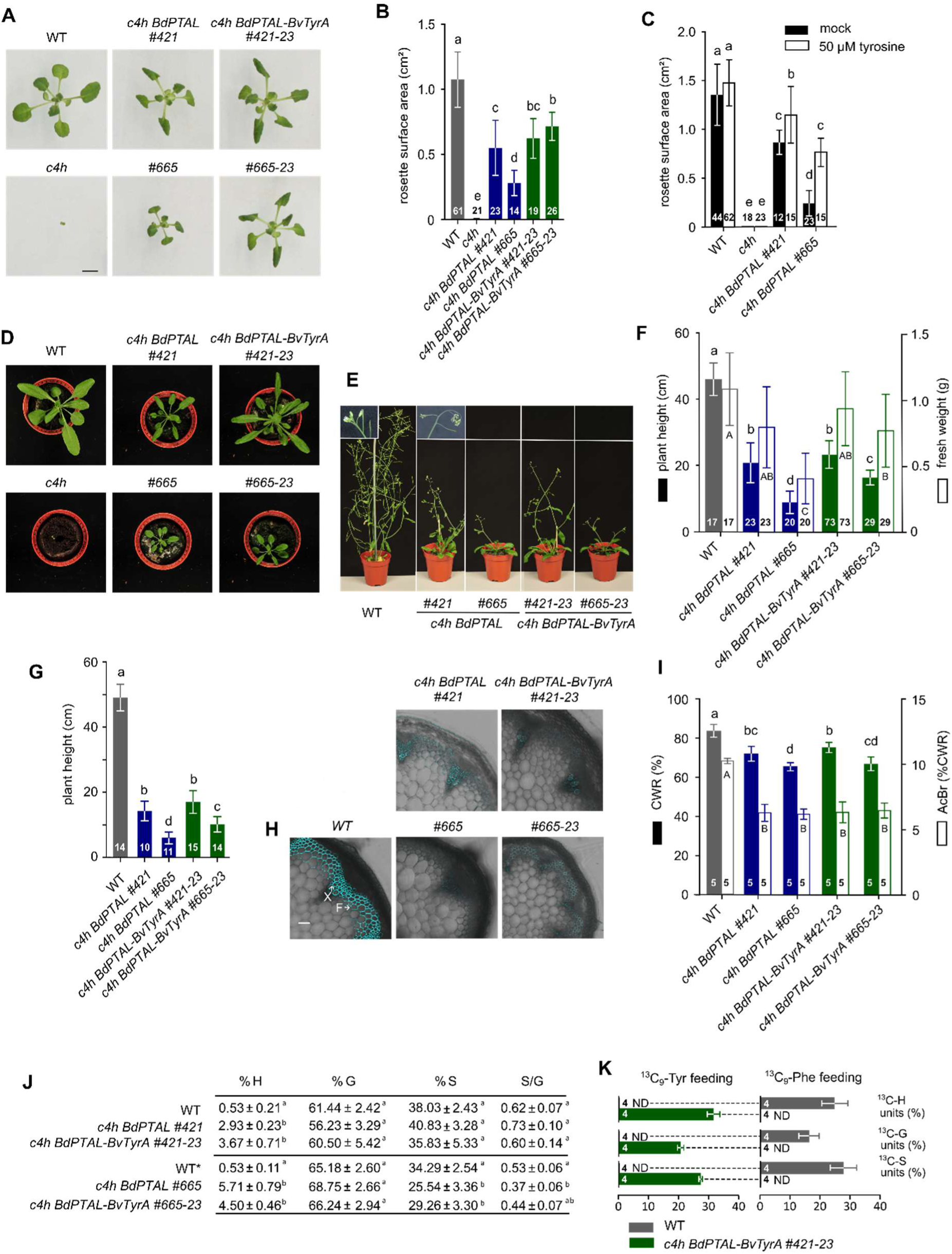
Establishing a tyrosine-based phenylpropanoid pathway in Arabidopsis. (**A**) Rosettes of *in vitro* grown WT, *c4h,* and *c4h*-complemented lines 21 DAS. Bar = 1 cm. (**B**) Projected rosette area of *in vitro* grown WT, *c4h* and *c4h*-complemented plants 21 DAS. (**C**) Projected rosette area of *in vitro* grown WT, *c4h* and *BdPTAL* overexpression lines 21 DAS, growing on 0.5xMS medium supplemented with 50 μM tyrosine or equal volume DMSO (mock). (**D**) Rosettes of soil-grown WT, *c4h* and *c4h*-complemented lines four weeks after sowing. Pot diameter is 5.5 cm. (**E**) Inflorescence stems of 7-week-old soil-grown WT and *c4h*-complemente*d* lines. Pot diameter is 5.5 cm. Insets at the top display flowers and developing siliques of 8-week-old WT and *c4h BdPTAL #421* plants. Bar = 1 mm. (**F**) Plant height and biomass of above-ground plant tissue (rosette + stem) of soil grown WT and *c4h*-complemented lines 7 weeks after sowing. (**G**) Plant height of fully senescent WT and *c4h*-complemented lines. (**H**) Lignin autofluorescence in the inflorescence stem of Arabidopsis WT and *c4h*-complemented lines. Bar = 50 µm. F: interfascicular fibers, X: xylem. (**I**) CWR and lignin content of the lower part of the fully senesced stem of WT and *BdPTAL* overexpression lines. The CWR determined gravimetrically after sequential extraction and expressed as a percentage of dry weight. The lignin content was assayed by the AcBr method and expressed as a percentage of CWR. (**J**) Lignin composition of the CWR of WT and *c4h*-complemented lines determined with thioacidolysis followed by GC-MS analysis. The relative proportions of the different lignin units were calculated based on the total thioacidolysis yield. S/G was calculated based on the absolute values for S and G. Values are given as average ± statistical deviation (n=5). Thioacidolysis for *c4h BdPTAL-BvTyrA #665* and *c4h BdPTAL-BvTyrA #665-23* was performed as an independent experiment. *WT control for thioacidolysis comparing WT to *c4h BdPTAL* #665 and *c4h BdPTAL-BvTyrA* #665-23. (**K**) Proportion of labeled precursors incorporated into each of the three main lignin units (H, G, and S) in 21-day-old *in vitro* grown plants grown on MS medium with isotopically labeled phenylalanine or tyrosine. The relative proportions of the different lignin units were calculated based on the total thioacidolysis yield (100% is the total area under the curve; n=4). Statistical significance was determined using an Anova test followed by a Tukey posthoc test. Numbers in the bars represent the number of repeats. Equal letters represent no statistical difference at the 0.05 significance level. Error bars represent standard deviation. AcBr: acetylbromide / BdPTAL: *Brachypodium distachyon* PHENYLALANINE TYROSINE AMMONIA LYASE / BvTyrA: *Brachypodium distachyon* AROGENATE DEHYDROGENASE / C4H: CINNAMATE-4-HYDROXYLASE / CWR: cell wall residue / PTAL: PHENYLALANINE TYROSINE AMMONIA LYASE / DAS: days after stratification / Tyr: tyrosine / WT: wild type

The partial growth recovery observed *in vitro* was also visible in four week-old plants grown in soil (**Fig. 2D**). By seven weeks, all restored lines developed inflorescence stems; however, these were considerably shorter compared to WT plants and exhibited apical desiccation (**Fig. 2E**). Quantitative analysis of total biomass, including both rosette and stem tissues, corroborated the *in vitro* findings. Among the restored lines, *c4h BdPTAL #421* showed a more robust phenotypic rescue relative to *c4h BdPTAL #665*, and the *BdTyrA* construct exerted its most pronounced additive effect in the *c4h BdPTAL #665* background (**Fig. 2F**). At senescence, the *c4h BdPTAL #665* line reached a final height of 6.0 cm, and consistent with the rosette surface and biomass data, the *c4h BdPTAL* #421 line performed significantly better, reaching a final height of 14.2 cm (**Fig. 2G**). The overexpression with *BvTyrA* in *c4h BdPTAL #421* did not result in any further increase in plant height. Despite the remarkable growth restoration in the complemented lines compared to the *c4h* mutant, which never formed an inflorescence stem, these plants remained much shorter than WT plants, which had an average height of 48.9 cm.

The effect of engineering a dedicated Tyr-derived phenylpropanoid pathway on stem morphology and lignin deposition was analysed via cross-sections. The *c4h* mutant was excluded from this analysis, as it does not develop stems. WT plants displayed open, round vessels and clear lignin autofluorescence (**Fig. 2H**). In contrast, vessels of *c4h BdPTAL* plants were smaller, collapsed, and showed a clear reduction in lignin UV autofluorescence, whereas interfascicular fiber cell walls displayed even weaker signals, indicating a strong reduction in lignin deposition in both cell-types. *BvTyrA* overexpression in these lines had no major effect on lignin autofluorescence. Lignin characteristics of the inflorescence stems in different lines were subsequently determined using AcBr and thioacidolysis. All transgenic lines had significantly less CWR compared to WT (82.8%), with the lowest in *c4h BdPTAL* #*655* (64.9%), and the highest in *c4h BdPTAL-BvTyrA #421-23* (74.0%), but no differences were found between *c4h BdPTAL* and their corresponding *BvTyrA-*overexpression lines (**Fig. 2I**). The AcBr lignin content of the CWR was significantly reduced in all *c4h BdPTAL* and *c4h BdPTAL-BvTyrA* lines compared to WT but did not differ between transgenic lines (**Fig. 2I**). Interestingly, thioacidolysis GC-MS analysis revealed the lignin of the restored lines to be enriched in H units: these accounted for only 0.53% of the total amount of thioacidolysis-released lignin monomers in WT, but reached to levels between 2.93% (*c4h BdPTAL #421)* and 5.71% (*c4h BdPTAL #665)* in the restored lines (**Fig. 2J**). While the relative amount of G and S units were unchanged in the *c4h BdPTAL #421* overexpression line, the relative amount of S units decreased in the *c4h BdPTAL #665* overexpression lines from 34% in WT to 26% and 29% in *c4h BdPTAL #665* and *c4h BdPTAL-BVTYRA #665-23*, respectively. This shift in relative S content resulted in a significant reduction in the S/G ratio, from 0.53 in WT to 0.37 in *c4h BdPTAL #665*.

To confirm the monofunctionally of the TAL pathway in the restored *c4h* mutants we performed isotopic tracing using ^13^C_9_-labelled amino acid precursors. The results showed that lignin biosynthesis in WT plants started only from ^13^C_9_-Phe, whereas in *c4h BdPTAL-BvTyrA #421-23* lines, lignin biosynthesis proceeded exclusively from ^13^C_9_-Tyr (**Fig. 2K**).

Collectively, these results show that a monofunctional TAL pathway can be functionally integrated into Arabidopsis to partially rescue the seedling lethal phenotype of the *c4h* mutant; however, these plants display stunted growth, vascular irregularities, and reduced lignin deposition enriched in H units.

### Cinnamic acid accumulation in restored *c4h* mutants causes auxin-related phenotypes

While the restored lines produced inflorescence stems, they exhibited apical desiccation upon flowering (**Fig. 2E**). As a result, *c4h BdPTAL #665*, and *c4h BdPTAL-BvTyrA #665-23* failed to produce mature siliques and were sterile. In contrast, *c4h BdPTAL #421* and *c4h BdPTAL-BvTyrA #421-23* occasionally produced seeds that were lighter in color compared to WT seeds (**Fig. 3A**). The seeds produced by *c4h BdPTAL #421* were viable, however, the germinated plants developed severely stunted rosettes and remained in a vegetative stage. The *c4h BdPTAL-BvTyrA #421-23* line was the best-performing restored line, with approximately 10% of the plants producing seeds. These plants had on average 2-3 siliques, each containing ∼5 seeds. When progeny from homozygous *c4h BdPTAL-BvTyrA #421-23* were sown on soil, all seeds germinated. These findings demonstrate that, despite exhibiting developmental abnormalities and reduced fertility, plants engineered to rely exclusively on a TAL-based phenylpropanoid pathway are nevertheless capable of completing reproduction and achieving a full life cycle.

**Figure 3:**
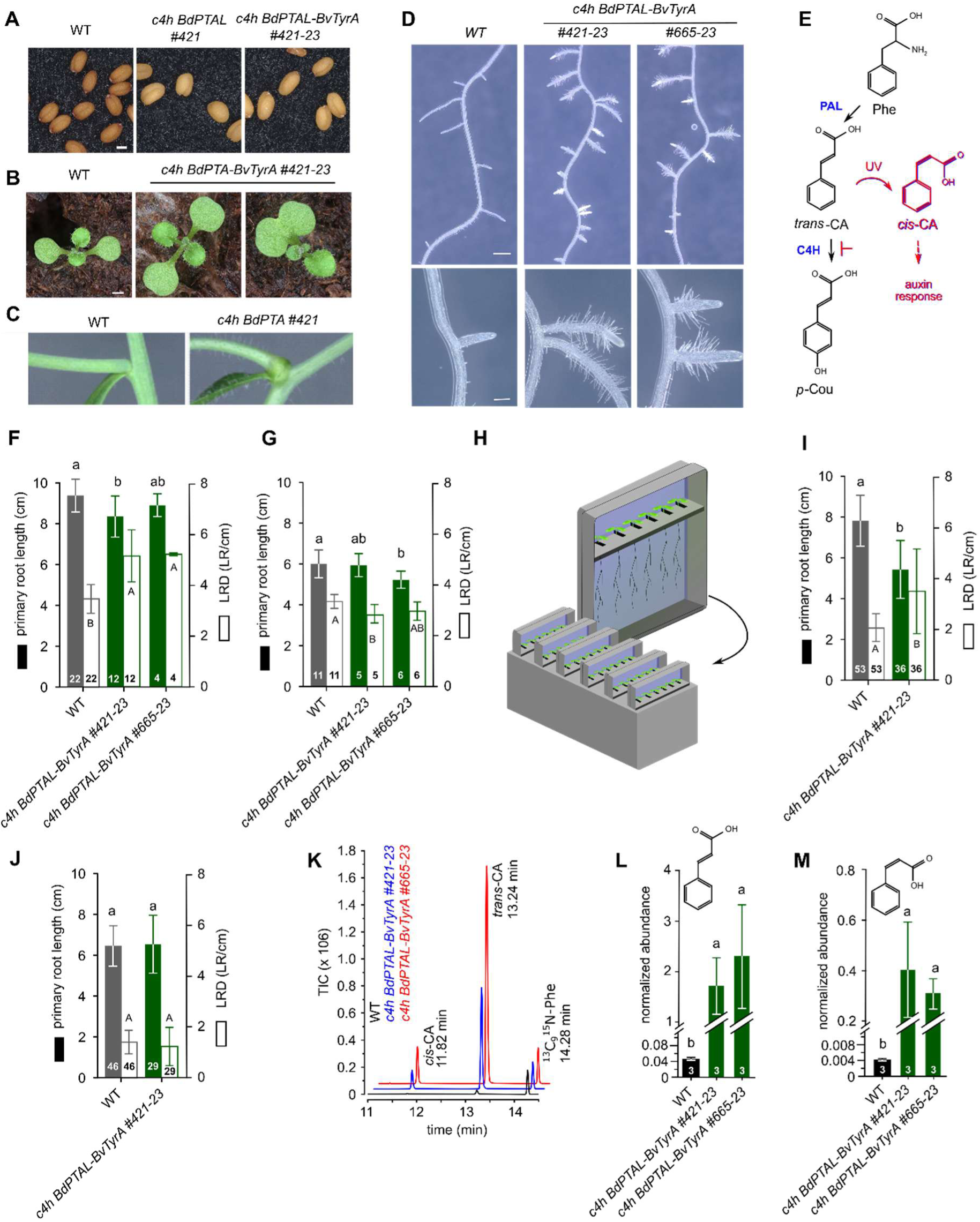
Metabolic engineering of the phenylpropanoid pathway modulates plant development. (**A**) Seeds of WT and homozygous *c4h* complemented plants. Bar = 20 µm. (**B**) Ten-day old progeny of WT and homozygous *c4h* complemented plants. Bar = 1 mm. (**C**) Stem branch junctions of 8-week-old WT and *c4h BdPTAL #421* plant. Bar = 1 mm. (**D**) Root phenotype of WT and homozygous *c4h* complemented plants 17 DAS. Top row: overview of the primary root with lateral roots. Bar = 1 mm. Bottom row: detail of the lateral roots with an increased number of root hairs. Bar = 100 µm. (**E**) Early phenylpropanoid pathway. The presumed accumulation of the auxin efflux inhibitor *cis*-CA upon blocking the C4H step is indicated in red. (**F**) Primary root length and LRD of WT and *c4h* complemented lines, growing under UV-B containing fluorescent light 17 DAS. (**G**) Primary root length and LRD of WT and *c4h* complemented lines, growing under LED light conditions 12 DAS. (**H**) Illustration of the D-root system used for *in vitro* plant growth, designed to maintain root development in darkness while exposing only the shoot to light. The setup was adapted from the system described in Silva-Navas *et al*., 2015. (**I**) Primary root length and LRD of WT and *c4h* complemented lines, growing under UV-B containing fluorescent light 14 DAS. (**J**) Primary root length and LRD of WT and *c4h* complemented lines, growing in the D-root system under UV-B containing fluorescent light 14 DAS. (**K**) Chromatogram illustrating the presence of *trans*- and *cis*-CA in roots of WT and *c4h* complemented lines growing under UV-B containing fluorescent light 17 DAS. ^13^C ^15^N-phenylalanine was used as an internal standard. (**L**) Normalized abundance of *trans*-CA in roots of WT and *c4h* complemented lines growing under UV-B containing fluorescent light 17 DAS. (**M**) Normalized abundance of *cis*-CA in roots of WT and *c4h* complemented lines growing under UV-B containing fluorescent light 17 DAS. All plants were grown under UV-B emitting fluorescent lamps, except plants used in (**G**), which were grown under LED lights depleted from UV-B. Plants used for (**J**) were grown in a D-root system as illustrated in (**H**). Statistical significance was determined using an Anova test followed by a Tukey posthoc test. Numbers in the bars represent the number of repeats. Equal letters represent no statistical difference at the 0.05 significance level. Error bars represent standard deviation. BdPTAL: *Brachypodium distachyon* PHENYLALANINE TYROSINE AMMONIA LYASE / BvTyrA: *Brachypodium distachyon* AROGENATE DEHYDROGENASE / C4H: CINNAMATE-4-HYDROXYLASE / CA: cinnamic acid / DAS: days after stratification / D-root: Dark-root / LED: light-emitting diode / LRD: lateral root density / PAL: PHENYLALANINE AMMONIA LYASE / Phe: phenylalanine / PTAL: PHENYLALANINE TYROSINE AMMONIA LYASE / FL: fluorescent light / TIC: total ion count / Tyr: tyrosine / TyrA: AROGENATE DEHYDROGENASE / UV: ultraviolet light / WT: wild type

While the apical desiccation and reduced seed set could result from a loss of mechanical strength or impaired water transport caused by defective vascular lignification (Li et al. 2015; Schilmiller et al. 2009; Huang et al. 2010), other developmental defects observed in the restored lines could not be attributed to lignin defects. For example, ∼10% of the homozygous *c4h BdPTAL-BvTyrA #421-23* seedlings showed developmental defects such as fused cotyledons (**Fig. 3B**) and local swellings were observed at branch junctions of *c4h BdPTAL* #421 and *c4h BdPTAL-BvTyrA* #421-23 lines (**Fig. 3C)**, a phenotype previously reported in leaky *C4H*-deficient *ref3* mutants (Schilmiller et al. 2009). In addition, the rosette leaves of all restored lines exhibited a characteristic leaf curling phenotype (**Fig. 2A, D**), and their roots displayed increased formation of lateral roots and root hairs (**Fig. 3D**). Interestingly, similar phenotypes were described in plants defective in auxin homeostasis and some have been linked to the accumulation of the auxin transport inhibitor *cis*-CA (Schilmiller et al. 2009; Steenackers et al. 2016). A similar mechanism may operate in our lines with a monofunctional TAL pathway, as they still produce *trans*-CA, but cannot convert it to *p*-coumarate due to the loss of C4H (**Fig. 3E)**.

To further investigate whether *cis*-CA accumulation contributed to the observed growth phenotypes, the roots of *c4h BdPTAL-BvTyrA* plants were analysed after growth on vertical agar plates under standard fluorescent light conditions. At 17 DAS, the *c4h BdPTAL-BvTyrA #421-23* line had a shorter primary root compared to WT (**Fig. 3F**). While the *c4h BdPTAL-BvTyrA #665-23* line followed the same trend, the difference was not significant from WT plants (*p =* 0.75). In addition, both restored lines exhibited higher lateral root density (LRD) relative to the WT (48% and 51% for #421-23 and #665-23, respectively) (**Fig. 3F**). A similar experiment was performed using LED lights without UV-B to avoid the photoisomerization of the phenylpropanoid intermediate *trans*-CA to its bioactive isomer *cis*-CA (Steenackers et al. 2016; Steenackers et al. 2019) (**Fig. 3E)**. While the *c4h BdPTAL-BvTyrA* plants grown under LED light still showed a small decrease in primary root length, although only significant for line #665-23, the strong increase in LRD was completely lost in both lines (**Fig. 3G**).

To evaluate the effects of potential *cis*-CA accumulation on root growth in *c4h BdPTAL-BvTyrA* plants, an adapted version of the D-root system was used (**Fig. 3H**), which exposes shoots of *in vitro* grown Arabidopsis seedlings to a normal light regime while shielding the lower portion of the plate to keep the roots in darkness (Silva-Navas et al. 2015). At 16 DAS, roots of *c4h BdPTAL-BvTyrA* #421-23 plants exposed to fluorescent light showed the expected reduction in primary root length (30.8%) and increase in LRD (72.3%) compared to roots of WT plants (**Fig. 3I**). However, when roots were shielded from light by growing the plants in the D-root system, no differences in primary root length and LRD were observed between roots of *c4h BdPTAL-BvTyrA* #421-23 and WT plants **(Fig. 3J)**. Besides demonstrating the involvement of light to increase LRD in the *c4h* restored line, the results obtained with the D-root system also showed that *cis*-CA cannot be transported from the illuminated shoot to the root system.

To unequivocally demonstrate the accumulation of *cis*-CA in *c4h* restored plants exposed to UV-B light, we used a previously optimized GC-MS protocol (El Houari et al. 2021). Using this method, both *cis*- and *trans*-CA were detected in roots of 17 days old WT plants grown under standard fluorescent light conditions, with *cis*-CA levels approximately 9.7% of those of *trans*-CA (**Fig. 3K)**. Both *c4h BdPTAL-BvTyrA* lines accumulated significantly higher levels of *trans*-CA, corresponding to a 40- and 54-fold increase compared to WT for line #421-23 and #665-23, respectively (**Fig. 3L)**. The accumulation of *cis*-CA was even more pronounced, with levels being 103- and 79-fold higher in *c4h BdPTAL-BvTyrA* lines #421-23 and #665-23 compared to WT plants (**Fig. 3M)**. Together, these results show that at least some of the auxin-related developmental defects observed in plants with an engineered monofunctional TAL pathway may arise from the accumulation of *trans*-CA, which can be converted to the auxin efflux blocker *cis*-CA under UV-B light conditions (Steenackers et al. 2016; El Houari et al. 2021).

## DISCUSSION

The phenylpropanoid pathway is a central metabolic pathway in plants that produces a wide array of phenolic compounds, including the precursors required for lignin biosynthesis. Because lignin is the second most abundant plant polymer after cellulose, substantial amounts of photosynthetically fixed carbon are routed into this pathway during periods of active lignification, underscoring the critical role of this pathway in plant carbon allocation and secondary metabolism (Boerjan, Ralph, and Baucher 2003).

While Phe serves as the primary precursor of the phenylpropanoid pathway in plants, Tyr offers a more direct biochemical entry point by circumventing both the hydroxyl group removal inherent to Phe synthesis and the subsequent 4-hydroxylation catalysed by the P450 enzyme C4H (Barros et al. 2016). Despite the lack of a Tyr specific pathway in plants, grasses have a bifunctional Phe/Tyr ammonia-lyase (PTAL) enzyme, which possesses the ability to use both Phe and Tyr as entry point for the phenylpropanoid pathway (Feduraev et al. 2020; Barros et al. 2016). Here, we demonstrated that a similar dual pathway can be engineered in Arabidopsis via the overexpression of *Brachypodium distachyon PTAL*, without detrimental phenotypic effects, illustrating the feasibility of non-grass species to use Tyr as entry for the phenylpropanoid pathway.

At the sequence level, the emergence of PTAL activity appears relatively easy, as a single amino acid substitution in the active-site is sufficient to convert Arabidopsis PAL1 into a functional PTAL (Watts et al. 2006), and just 2 amino acid switches are enough to convert the *Joinvillea ascendens* PAL into a PTAL (Takeda-Kimura et al. 2024). In addition, most plant genomes contain multiple *PAL* genes (i.e. Arabidopsis has four *PAL*s), providing the genetic redundancy required for neofunctionalization (Huang et al. 2010). Indeed, the emergence of PTAL activity in grasses has been attributed to a spontaneous mutation event (Peng et al. 2022). Intriguingly, although the biochemical modification required to generate a Tyr-active ammonia lyase seems fairly simple, analogous events have not been observed in plant lineages other than grasses from the Poaceae family, suggesting that a substantial evolutionary barrier prevents the conversion of PALs into PTALs (Barros and Dixon 2020). Furthermore, the subsequent transition from bifunctional PTAL to monofunctional TAL catalysing a Tyr-specific phenylpropanoid pathway also appears to be evolutionarily constrained in plants, as no evidence suggests that such conversion has ever occurred.

One hypothesis for why an exclusively Tyr-derived pathway has not evolved in plants is that intracellular Tyr availability may be insufficient to sustain such a demanding downstream metabolic route. Being an essential primary metabolite, diverting substantial amounts of Tyr into the phenylpropanoid pathway could compromise fundamental cellular processes such as protein synthesis. Consequently, the metabolic burden imposed by extensive Tyr withdrawal may create a selective pressure that limits the evolutionary feasibility of establishing a fully Tyr-dependent biosynthetic pathway. Interestingly, plants that possess PTALs, such as grasses, compensate for the increased demand for Tyr by producing it at levels more than tenfold higher than those observed in dicots like Arabidopsis (El-Azaz et al. 2023). This elevated Tyr production is attributed to the deregulation of key entry and exit points in the aromatic amino acid biosynthetic pathway (El-Azaz, Moore, and Maeda 2025; El-Azaz et al. 2023). Similarly, members of the Caryophyllales, including red beets, have feedback insensitive TyrA enzymes to allow the abundant production of Tyr-derived betalaines (Lopez-Nieves et al. 2022). Phylogenetic analyses in both grasses and beets have shown that the deregulation of TyrA enzymes predates the evolution of the downstream metabolic steps required for lignin and betalain biosynthesis (Lopez-Nieves et al. 2018; El-Azaz, Moore, and Maeda 2025). Together with observations that elevated Tyr levels can negatively affect plant growth in transgenic lines (de Oliveira et al. 2019), these findings support the hypothesis that, increased Tyr accumulation resulting from TyrA deregulation may have activated detoxification mechanisms, allowing the evolution of downstream sinks. For example, in *Beta vulgaris*, this likely contributed to the evolution of betalain biosynthesis, whereas in grasses it may have promoted the emergence of the Tyr-derived branch of the phenylpropanoid pathway. While this evolutionary based model cannot be dismissed, the absence of detrimental phenotypes in the PTAL overexpression lines do not support the need for elevated endogenous Tyr levels prior to the appearance of (P)TAL. Additionally, this Tyr-depletion hypothesis runs short in explaining why the pathway never developed towards a dedicated Tyr phenylpropanoid pathway.

An alternative theory to explain the absence of a Tyr-specific phenylpropanoid pathway in plants builds on the crucial role of *trans*-CA as an evolutionary driver to retain the Phe branch of the pathway (Maeda 2016). A Tyr-specific pathway bypasses the production of *trans*-CA, implying that plants with such pathway would lack this metabolite. Besides being a key intermediate of the canonical phenylpropanoid pathway, *trans*-CA also serves as a precursor to a variety of specialized metabolic pathways that contribute to the biosynthesis of ecologically and physiologically important compounds. For instance, *trans*-CA is a biosynthetic precursor to salicylic acid, a key phytohormone involved in systemic acquired resistance and defence signalling (Zhu et al. 2025). However, salicylic acid can also be produced independently from *trans*-CA via the isochorismate pathway (Hong et al. 2025). Additionally, certain hydroxycoumarins and stilbenes, compounds recognized for their antimicrobial and allelopathic activities, are also derived from *trans*-CA (Stringlis, De Jonge, and Pieterse 2019; Al-Khayri et al. 2023). Yet, these metabolites are synthesised via *p*-coumarate, which remains available through the Tyr-derived pathway, implying that the absence of *trans-*CA could influence but not deplete their formation.

In addition to its role as a pathway intermediate, the *cis*-isomer of CA can act as a signalling molecule, influencing auxin transport and modulating plant growth responses (Steenackers et al. 2016). The latter was clearly illustrated in our *c4h* restored lines. As a consequence of the metabolic engineering strategy, these lines produced *trans*-CA that build up due to the absence of C4H to channel it into the phenylpropanoid pathway (**Fig. 3L, M**). When grown under canonical UV-B emitting fluorescent lamps, these plants display classic auxin-related phenotypes such as enhanced lateral root formation and increased root hair density (**Fig. 3D**). The association between these phenotypes and elevated *cis*-CA levels, together with the attenuation of the phenotypes under UV-depleted LED light or in dark grown roots, suggests that in the *c4h* restored lines, accumulated *trans*-CA undergoes photoisomerization to its *cis*-isomer, that can then act as an inhibitor of auxin transport (Steenackers et al., 2016). In addition, we identified *cis*-CA in WT seedlings, implying its participation in regulatory mechanisms underlying plant development. As plants can grow in the absence of UV without detrimental defects (e.g. our LED light conditions), the light-mediated isomerization of CA is likely not crucial for plant development. However, this does not preclude a potential role of the *cis*-isomer in plant physiology as the bioactive molecule still could be produced through alternative mechanisms, such as by enzymatic conversion. In fact, *cis*-CA has been detected in Arabidopsis seedlings grown in the absence of UV-light, supporting the existence of such CA isomerase (El Houari et al. 2021).

While the PTAL-engineered lines described in this paper provide a valuable tool for elucidating the physiological functions of *trans*- and *cis*-CA in greater detail, they fall short in addressing the fundamental question of whether CA constitutes an essential metabolite in plants, that hinders the emergence of a Tyr-specific phenylpropanoid pathway (Maeda 2016). A more definitive assessment of CA’s essentiality would require experimental approaches that directly test the consequences of its complete loss under various physiological conditions. Theoretically, plants unable to produce *trans*-CA can be obtained by blocking the phenylpropanoid pathway at its first step (i.e. the deamination of Phe) and bypassing the inactivated conversion with a Tyr-specific pathway. However, in contrast to *C4H* used in this study and which is a single copy gene in Arabidopsis, the *PAL* gene family is represented by four members, complicating metabolic engineering. Notably, quadruple *PAL* T-DNA insertion mutants are described, but these still show residual PAL activity and will consequently still produce *trans*-CA, thereby compromising their suitability for the intended investigation into the critical role of this metabolite (Huang et al. 2010). In addition, the proposed strategy cannot rely on the *PTAL* construct we used in this study to bypass PAL, as the PAL activity of this bifunctional enzyme would mitigate the inactivation of the PAL step, and still produce *trans*-CA. Alternatively, a monofunctional TAL enzyme derived from bacteria could be employed (Jendresen et al. 2015). For example, expression of *Flavobacterium johnsoniae* TAL in Arabidopsis resulted in elevated levels of flavonoids and anthocyanins relative to wild-type controls (Nishiyama et al. 2010). However, our preliminary complementation experiments using a panel of bacterial TALs showed that these monofunctional enzymes do not function sufficiently *in planta* to rescue the *c4h* mutant (Van Beirs *et al*, in preparation). As no monofunctional TALs with adequate activity are currently available, and so far no monofunctional TALs have been engineered from bifunctional PTALs, generating plants completely lacking CA is not possible with the tools presently at hand.

In conclusion, this study demonstrates the potential for engineering both a dual Phe/Tyr-derived phenylpropanoid pathway as well as an exclusive Tyr-specific pathway in Arabidopsis, despite the latter not occurring naturally in plants. Our findings suggest that CA may act as a constraining factor in the evolutionary establishment of a Tyr specific biosynthetic pathway. Furthermore, the genetic evidence presented here supports a biological role for *cis*-CA in plants, and that this molecule may act as a UV receptor molecule shaping plant growth and development by altering auxin homeostasis.

## MATERIALS & METHODS

### Plant materials, chemicals and growth conditions

*Arabidopsis thaliana* lines of the Columbia 0 (Col 0) ecotype were used for all assays. The *C4H* T-DNA insertion line (GABIKAT GK-753B06; (Kleinboelting et al. 2012) was obtained from the Nottingham Arabidopsis Stock Centre (NASC), and the Arabidopsis line overexpressing the *Beta vulgaris p35S::BvTyrA* construct was received from Hiroshi Maeda (de Oliveira et al. 2019). Before sowing, seeds were vapor-phase sterilized by incubating them in open 1.5 mL-Eppendorf tubes overnight in a desiccator containing chlorine fumes generated by the reaction of 3 mL concentrated HCl with 100 mL commercial bleach. Surface-sterilized seeds were sown on either square (12 × 12 cm) or round (16 cm diameter). Petri dishes contained 0.8% (w/v) agar-solidified half-strength Murashige and Skoog (MS) medium (pH 5.7). The medium composition per litre included 2.15 g MS basal salt mixture powder (Duchefa), 10 g sucrose, and 0.5 g 2-(N-morpholino)ethanesulfonic acid (MES) monohydrate (Duchefa). For chemical complementation, the medium was supplemented with tyrosine (Sigma) dissolved in DMSO (Sigma; stock concentration 40 mM; final concentration 100 µM). After sowing, plates were incubated at 4 °C for two days and subsequently positioned either vertically or horizontally under long-day conditions (16 h light/8 h dark) at 21 °C. Plants grown in soil were maintained under identical long-day photoperiod conditions using Jiffy-7 peat pellets, which had been pre-soaked in water. When not mentioned otherwise, light was provided by full-spectrum Osram L 58W/840 Lumilux Cool White fluorescent lamps (FL) delivering a photosynthetic photon flux density (PPFD) of 90 µmol m⁻² s⁻¹ at shelf level, measured using a SpectroSense2+ quantum sensor (Skye Instruments Ltd, UK). The Osram lamps emitted 0.03 μmol m^−2^ s^−1^ UV-B light measured at shelf level by the SpectroSense2+ meter equipped with an SKU 430/SS2 UV-B sensor. When LED lighting conditions are referenced, Valoya LED tube L35-NS12 units were used, providing a PPFD of 90 µmol m⁻² s⁻¹ at shelf level. The spectral output exhibited three principal peaks at approximately 400, 450, and 620 nm, and the tubes did not emit UV-B radiation.

### Vector construction and generation of transgenic plant lines

A 2-kb genomic region upstream of the *C4H* (AT2G30490) ATG start codon was considered the *C4H*-promoter region and was PCR-amplified from *A. thaliana* Col 0 genomic DNA using Phusion^TM^ High-Fidelity DNA polymerase (Thermo Fisher). The *Brachypodium distachyon PTAL* (XP_003575396) coding sequence (*BdPTAL* CDS) was PCR amplified using Q5^®^ High-fidelity polymerase (New England Biolabs) using a CDS-containing plasmid as template (Barros et al. 2016). The primers used for the PCR reactions contained BsaI restriction sites and unique 4-bp overhangs required for Golden Gate cloning (Supplementary Table S2;(Lampropoulos et al. 2013)). The amplification products (i.e. *AtC4H* promotor and *BdPTAL* CDS) were gel-purified (GeneJET Gel Extraction Kit, Thermofisher) and cloned into the pGGA000 (Vector ID 7_43) or pGGC000 (Vector ID 7_45) Golden Gate entry vector using T4 Ligase (New England Biolabs). The ligation mixture was subsequently used to transform *Escherichia coli* DH5α competent cells via heat shock at 42 °C for 45 seconds, followed by recovery in 0,5 mL LB medium at 37 °C for 1 hour with shaking at 350 rpm in an Eppendorf ThermoMixer F2.0. Transformed cells were plated on LB agar supplemented with the appropriate antibiotic for selection and incubated overnight at 37 °C. Individual colonies were screened by colony PCR using vector- and insert-specific primers to confirm the presence of the different building blocks. Positive clones were further validated by plasmid extraction and sequencing (Eurofins). After sequence confirmation, both entry vectors were combined with a His6x-tag containing entry vector into the Golden Gate pGGK-AG destination vector, which resulted in the p*C4H*::*BdPTAL* expression clone. The selection of the correct expression clone was performed in *E. coli* DH5α as described above, and the sequence-confirmed expression clone was introduced into *Agrobacterium tumefaciens* strain C58C1 by electroporation (Bio-Rad 0.2 cm electrode gap cuvette, 2.5 kV with a Bio-Rad MicroPulser electroporator). The obtained Agrobacterium strain was used to transform heterozygous *C4H/c4h* plants using the floral dip method (Clough and Bent 1998). T1 transgenic plants were selected on ½ × MS medium containing 50 µg/ml kanamycin. To track the segregation of the T-DNA tagged *C4H* allele, Edwards-extracted genomic DNA of individual seedlings was used for PCR-based genotyping(Edwards, Johnstone, and Thompson 1991) using one primer pair flanking the T-DNA insertion site to detect the wild-type allele, and a second pair consisting of a gene-specific primer and a T-DNA border primer to detect the insertion allele (Supplementary Table S2). Over the course of three plant generations, two independent single locus lines were selected (*C4H/c4h BdPTAL* lines #421 and #665). Both *C4H/c4h BdPTAL* lines were crossed with the *p35S::BvTyrA* #23 line. Plants homozygous for *p35S*::*BvTyrA* and *pC4H*::*BdPTAL*, but heterozygous for *C4H/c4h* were obtained through selection on gentamycin (50 μg/mL; *p35S*::*BvTyrA* selection), kanamycin (50 μg/mL; *pC4H*::*BdPTAL* selection), and sulfadiazine (25 mg/L; *C4H/c4h* selection) over the course of three additional plant generations. The genotype of the plants of the different progenies was confirmed by PCR using gene and allele specific primers (Supplementary Table S2).

### Quantitative Real-Time PCR

Arabidopsis plants were grown on ½ × MS medium as described above. At 2 days after stratification (DAS), 5 seedlings from each line were collected, flash-frozen in liquid nitrogen, and homogenized for further analysis (Retsch MM200 mixer mill). For each line, three biological replicates were used. RNA was isolated using ReliaprepTM RNA tissue miniprep kit (Promega) followed by a DNase treatment. RNA concentration and purity were assessed using an ND-1000 NanoDrop spectrophotometer (Thermo Fisher Scientific). For cDNA synthesis, 1 µg of total RNA was reverse transcribed with 4 µL of qScript® cDNA SuperMix (Quantabio) in a 20 µL reaction containing RNase/DNase-free water. Each biological repeat was analyzed in three technical repeats with 384-multiwells plates. Relative gene expression was quantified using the LightCycler 480 II apparatus (Roche) with the SYBR Green I Master mix Kit (Roche Diagnostics) according to the manufacturer’s instructions. The expression of the selected genes was determined with the Roche LightCycler 480 combined with the SYBR Green I master Kit (Roche Diagnostics) using the following PCR protocol: one activation cycle of 10 min (95 °C); 45 amplification cycles of 10 s (95 °C), 10 s (60 °C) and 10 s (72 °C). Each biological repeat sample was run in triplicate to identify potential technical variation that could have occurred during pipetting. The cDNA was diluted 10 times prior to use. Every sample had a total volume of 20 μL, consisting of 10 μL 2× SYBR Green mix, 8 μL primer mix (1 μM), and 2 μL diluted cDNA. Fluorescence values were exported from the Lightcycler 480 program, after which Ct values, normalization factors, and primer efficiencies were calculated according to (Ramakers et al. 2003) using two reference genes: UNIQUITIN-CONJUGATING ENZYME 21 (UBC21; AT5G25760). Primers are listed in Supplementary Table S2.

### Projected rosette area and total leaf area

The quantification of the projected rosette area was performed on Arabidopsis plants grown horizontally on ½ × MS medium in round Petri dishes (diameter 16 cm). At 21 DAS, plates were scanned using the Scanmaker 9800XL, and the projected rosette area of individual plants was measured using ImageJ software(Schneider, Rasband, and Eliceiri 2012). Data were analysed and visualized using GraphPad Prism10 software.

### Histochemistry

Arabidopsis plants were grown on soil as described above. Seven weeks after sowing, the primary inflorescence stem was harvested by cutting the stem just above the rosette. The bottom 1 cm of the main stem was removed, and the subsequent 1 cm was embedded in 7% (w/v) agarose. The agarose plugs were sliced in 100 μm sections using a 5100mz-vibratome (Campden Instruments). Lignin autofluorescence was imaged using the Zeiss LSM 780 microscope with a *Plan*-Apochromat 103 (0.45 M27) objective. The fluorescence signal for lignin was obtained using an excitation wavelength of 350 nm and an emission wavelength range of 407 to 479 nm.

### Lignin cell wall analysis

For analysis of the lignin content and composition, the bottom 10 cm (excluding the bottom 1 cm) of the senesced primary inflorescence stem was chopped into 2 mm-pieces, and samples were pooled per two individuals. Preparation of cell wall residue and acetyl bromide extraction were performed as previously described in (Van Acker et al. 2013). The general lignin composition was determined using thioacidolysis as essentially described by Robinson and Mansfield (2009a) and modified by De Meester et al. (2022). For sample derivatization, 500 µL of the organic phase of every sample was evaporated to dryness under vacuum, and if needed stored at −20 °C and resuspended in 200 µL of dichloromethane. Samples were derivatized by combining 10 µL of resuspended sample with 20 µL of pyridine and 100 µL of *N*,*O*-bis(trimethylsilyl)acetamide by incubation (750 rpm) for 2 h at 25 °C. The monomers released upon thioacidolysis were detected with gas chromatography (GC)_ as their trimethylsilyl ether derivatives on a 7890B GC system (Agilent, Santa Clara, CA, USA) coupled with a 7250 GC/QTOF mass detector (Agilent, Santa Clara, CA, USA, Supplemental Table S3). A 1-µL aliquot was injected in splitless mode on a VF-5ms capillary column (40 m x 0.25 mm x 0.25 µm; Varian CP9013; Agilent) at a constant helium flow of 1.2 mL/min. After injection, the oven was held at 130 °C for 3 min, ramped to 200 °C at a rate of 10 °C.min^-1^, ramped to 250 °C at a rate of 3 °C /min and held at 250 °C for 5 min, then ramped to 320 °C at a rate of 20 °C.min^-1^ and held at 320 °C for 5 min. The injector, mass spectrometry transfer line, the mass spectrometry ion source, and the quadrupole were set to 280 °C, 280 °C, 230 °C and 150 °C, respectively. The mass spectrometry detector was operated in EI mode at 70 eV. Full EI-mass spectrometry spectra were generated for each sample by scanning the *m*/*z* range of 50 to 800 with a solvent delay of 10 min. Quantitative evaluation was carried out with Masshunter Quantitative Analysis (version 10.0) based on the specific quantifier for each compound. Response factors for H, G, and S units were determined according to Yue et al. (2012) (Supplemental Table S3). Data were analysed and visualized using GraphPad Prism10 software.

### Isotopic Labeling Experiments

Arabidopsis plants were grown in 68 × 110 mm culture tubes (Greiner) containing 1x MS media with vitamins (Phytotech) at pH 5.8, supplemented with sucrose (3% w/v) and phytagel (Sigma-Aldrich). For isotopic labeling, the medium was additionally supplemented with 0.1 mM of ^13^C_9_-Phe or ^13^C_9_-Tyr (Sigma-Aldrich). Approximately 15-20 seeds were evenly distributed onto the medium in each container. After a 2-day stratification at 4 °C, the tubes were transferred to continuous light and maintained at 24 °C. Wild-type and mutant plants were grown in separate batches and harvested three weeks after germination. For each genotype, roots and shoots from all plantlets were collected, pooled, and stored at −80 °C until further analyses. To quantify the levels of ^13^C-labeled monolignols, plant tissues were ground in liquid N_2_, followed by cell wall extraction and overnight lyophilization. Lignin extraction was performed using thioacidolysis following established protocols ((Barros et al. 2016; Barros et al. 2022) and the peak areas of the primary lignin monomers H (239 *m*/*z*), G (269 *m*/*z*), S (299 *m*/*z*), were identified by GC-MS. Monolignol fragments lose two carbons off the side chain during GC-MS fragmentation after thioacidolysis, therefore peak areas with M0 and M+7 masses were collected for each monomer. To calculate the percentage of monolignol incorporation for each subunit, the labeled (M+7) peak area were divided by the unlabeled (M0) plus the labeled (M+7) peak areas and multiplied by 100. Data were analyzed and visualized using GraphPad Prism10 software.

### Detection of *cis*- and *trans*-cinnamic acid

Arabidopsis plants were grown on ½ × MS medium as described above. At 17 days after stratification (DAS), 3 seedlings from each line were collected, flash-frozen in liquid nitrogen, and homogenized for further analysis (Retsch MM200 mixer mill). For each line, three biological replicates were used. Metabolites were extracted with 1ml methanol for 15 min. Two hundred µL of the methanol extracts, containing 2,000 pg.µL^-1^ ^13^C_9_^15^N-phenylalanine (Sigma) as internal standard, were lyophilized using a Labconco CentriVap SpeedVac concentrator and subsequently derivatized via trimethylsilylation using 50 µL N-methyl-N-(trimethylsilyl)trifluoroacetamide (MSTFA; Sigma) with pyridine in a 5:1 ratio. The derivatization reaction was carried out at 37 °C for 60 minutes under continuous agitation to ensure complete reaction. Following derivatization, 1 µL of the samples was injected in splitless mode into an Agilent 8890 GC coupled to an Agilent 7010B Triple Quadrupole MS. The injector temperature was set at 280 °C. Chromatographic separation was achieved using an HP-5ms capillary column (30 m × 0.25 mm i.d., 0.25 μm; Agilent, 19091S-433UI) with helium as the carrier gas at a constant flow rate of 0.8 mL.min⁻¹. The oven temperature program was as follows: initial hold at 80 °C for 1 minute, ramp to 250 °C at 10 °C.min⁻¹, followed by a rapid increase to 320 °C at 50 °C.min⁻¹, with a final hold at 320 °C for 3 minutes. The mass spectrometry transfer line, ion source, and quadrupole temperatures were set to 280 °C, 230 °C, and 150 °C, respectively. Electron ionization (EI) was performed at 70 eV. For quantitative analysis, the MS was operated in multiple reaction mode (MRM). The specific MRM transitions for *cis*- and *trans*-cinnamic acid are provided in Supplementary Table S4. Relative quantification was based on the integration of the quantifier ions, normalized to the internal standard.

### *In vitro* dark-grown roots (D-root system)

The dark-grown root system was based on the D-root device introduced by (Silva-Navas et al. 2015). The boxes holding the square (12 × 12 cm) plates and the combs separating the light-exposed side from the root growth area in the plate were designed in Blender (www.blender.org), and the objects were printed with a Bambu Lab X1 Carbon 3D Printer using black polylactic acid and a 0,2 mm nozzle size. The STL files are provided as supplementary information. The boxes were designed in a way that multiple plates could be placed in a single box. The 3D-printed combs were slightly modified from the original design by replacing the open structure with a closed configuration featuring narrow slits, allowing seedlings to grow with their shoots emerging on one side and their roots extending on the opposite side (Fig 3J). The modified combs maximized the shielding of the root zone from the light and minimized the contact area of the medium between the light-exposed side and root growth area, reducing the diffusion of compounds between both regions. The experiment was initiated by spreading surface-sterilized seeds on ½ × MS medium. Following sowing, the seeds were incubated at 4 °C for two days and subsequently incubated under long-day growth conditions (16 h light/8 h dark) at 21 °C for an additional two days to trigger germination. Seedlings were transferred to 12 x 12 cm square plates, containing 25 mL MS-medium. Prior to solidification, a 3D-printed comb was inserted into the medium. Once the medium had solidified, seeds were carefully positioned along the upper edge of the comb, aligned with the open slits. The plates were closed, sealed with Urgopor and placed vertically into custom-designed boxes, which were inclined at a 10° angle and incubated under long-day growth conditions. Twelve days after transfer, the root growth was analysed by measuring the primary root length and the number of lateral roots. A second set of seeds, serving as a control, underwent an identical treatment protocol, with the exception that the plates were not placed in a D-root growth box.

## Acknowledgements

We thank Richard Dixon for providing the *BdPTAL* expression vector and Hiroshi Maeda for sharing the Arabidopsis *p35S::BvTyrA* overexpression line. Kristof Verleye is acknowledged for 3D-printing and Karel Spruyt for photographic work. We thank the VIB Bio-Imaging core for microscopy assistance, and Kaho Cheng for the assistance with the GC-MS chromatograms.

## SUPPLEMENTARY INFORMATION

**Figure S1:**
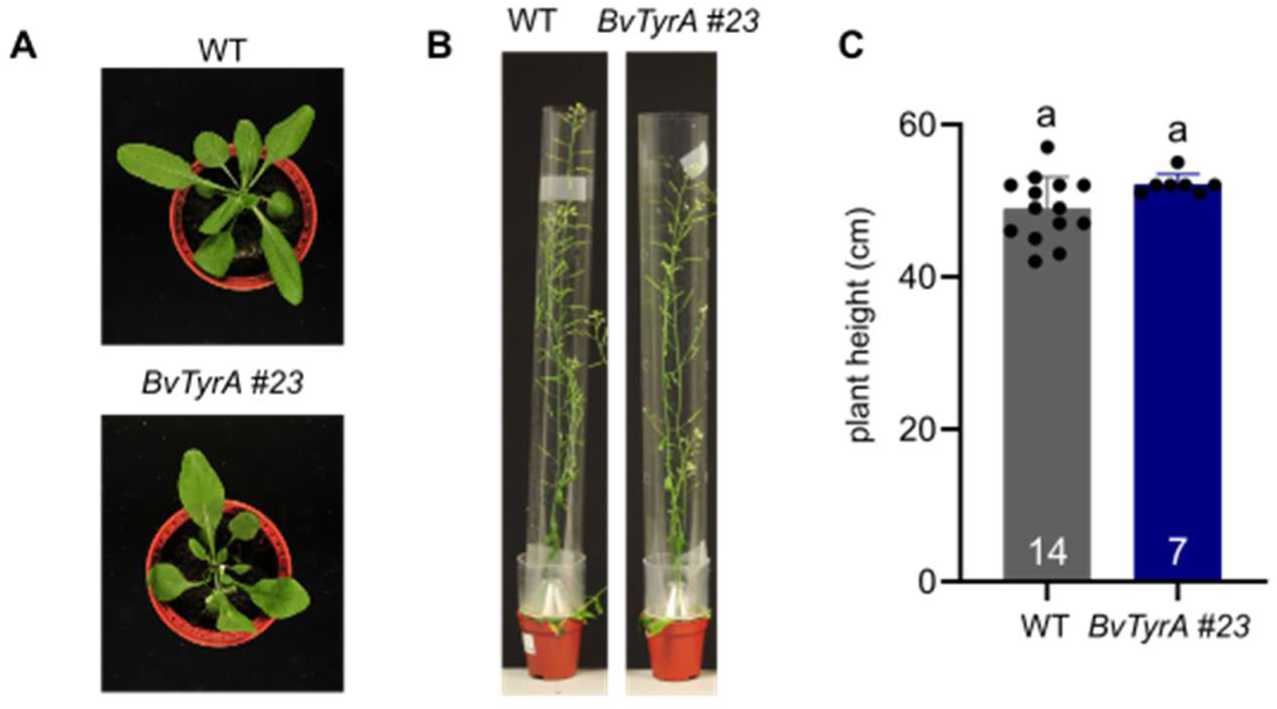
Phenotype of *BvTyrA* overexpression line #23. A) Rosettes of 4-week-old soil-grown WT and the *BvTyrA* overexpression line #23. Pot diameter is 5.5 cm. B) Inflorescence stems of 7-week-old WT and the *BvTyrA* overexpression line #23. Pot diameter is 5.5 cm. C) Plant height of fully senesced soil-grown WT and the *BvTyrA* overexpression line #23. Statistical significance was determined using an Anova test followed by a Tukey posthoc test. Numbers in the bars represent the number of repeats. Equal letters represent no statistical difference at the 0.05 significance level. Error bars represent standard deviation.

**Figure S2:**
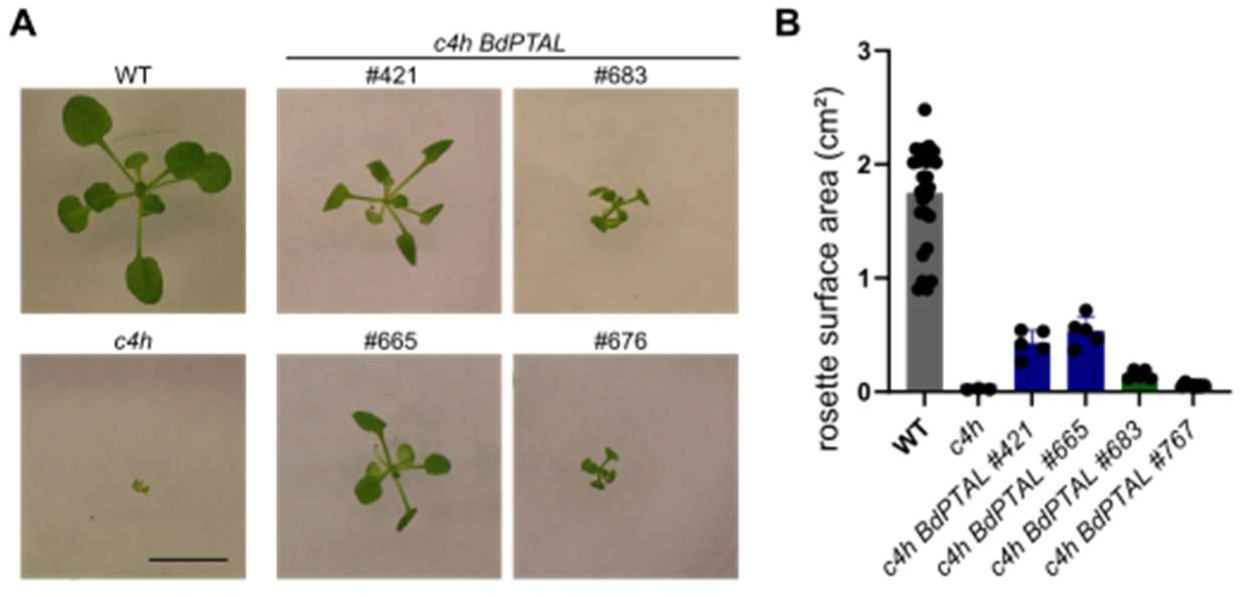
Phenotype of c4h-complemented lines #421 and #683. A) Rosettes of *in vitro* grown WT, *c4h,* and weak *c4h*-complemented lines #421 and #683, 21 DAS. Bar = 1 cm. B) Projected rosette area of *in vitro* grown WT, *c4h* and weak *c4h*-complemented lines #421 and #683, 21 DAS. Statistical significance was determined using an Anova test followed by a Tukey posthoc test. Numbers in the bars represent the number of repeats. Equal letters represent no statistical difference at the 0.05 significance level. Error bars represent standard deviation.

**Table S1:**
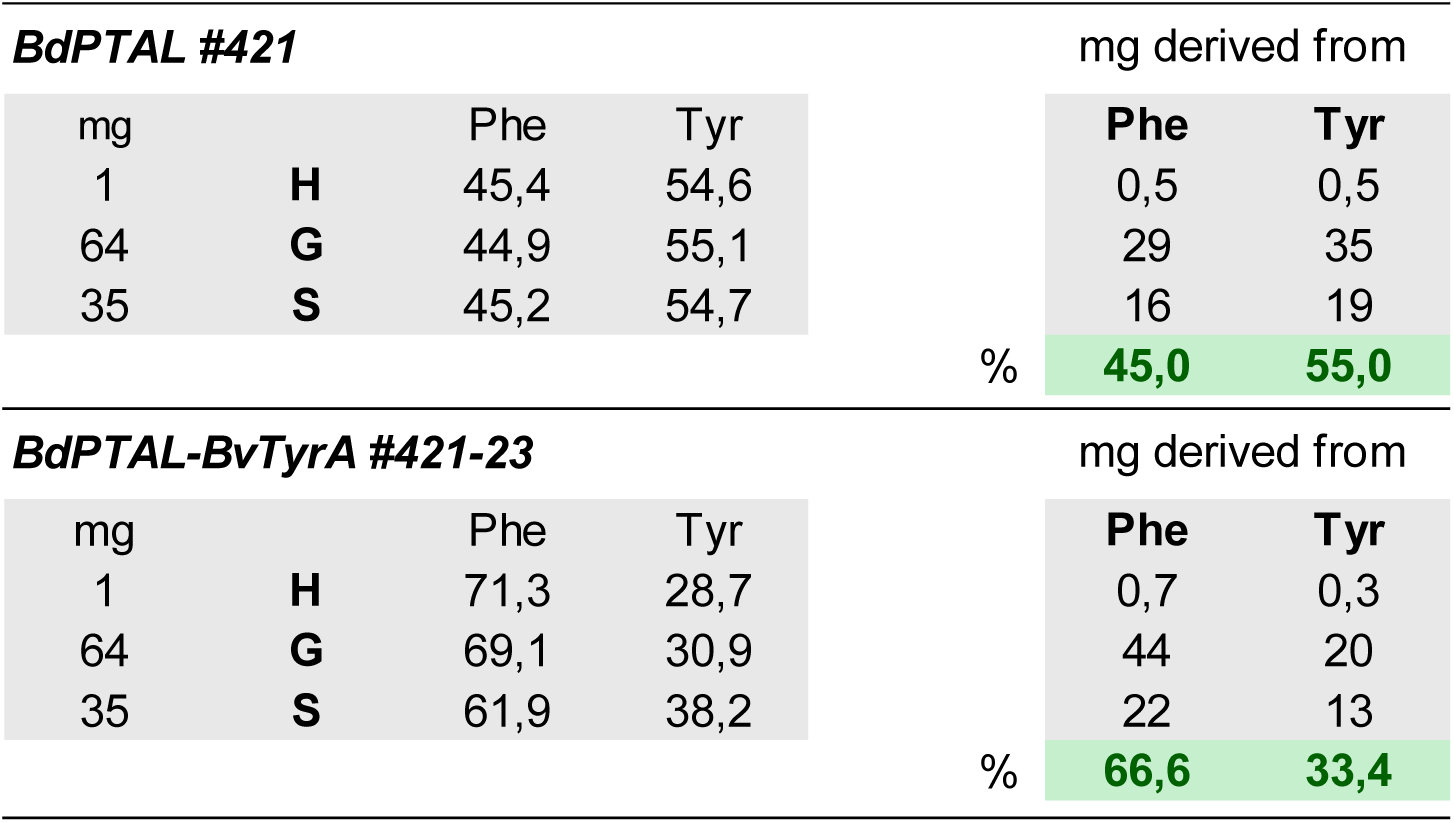
Incorporation of labeled precursors in lignin. Proportion of labeled precursors incorporated into each of the three main lignin units (H, G, and S) in 21-day-old in vitro grown plants grown on MS medium with isotopically labeled phenylalanine or tyrosine.

**Table S2:**
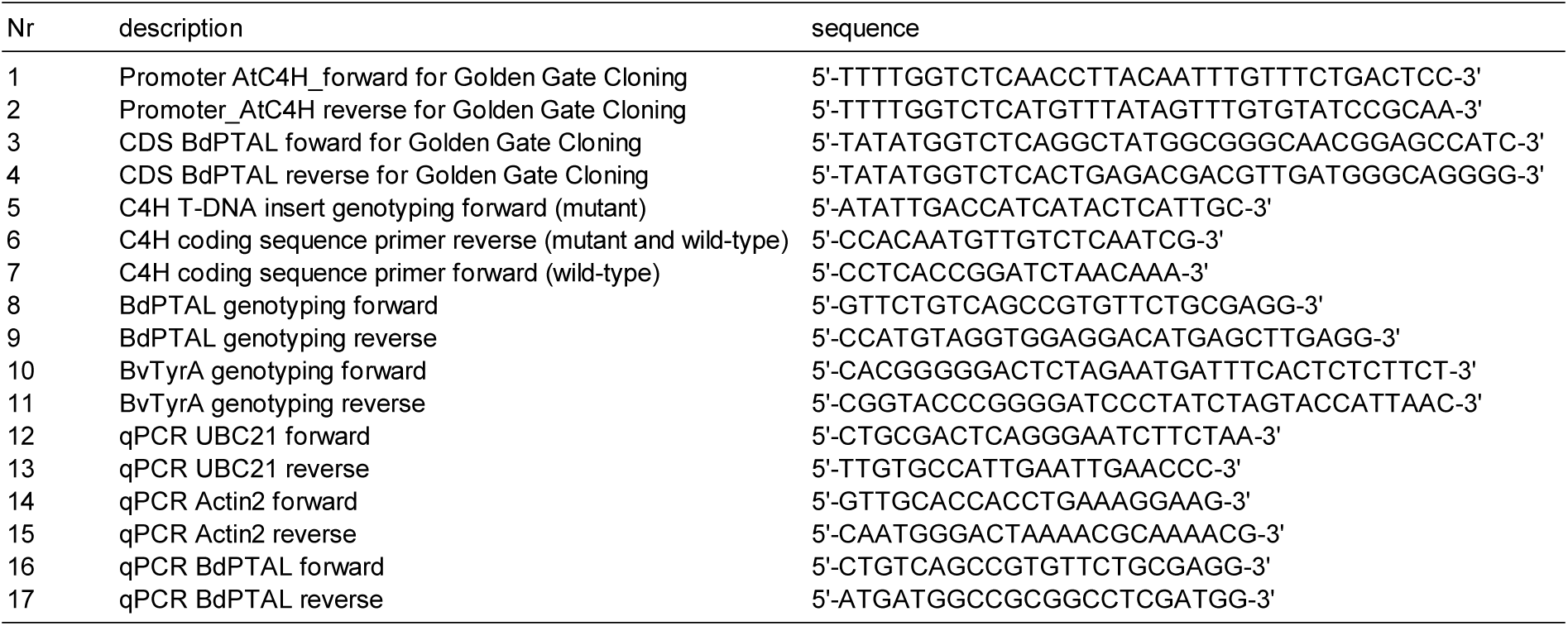
Oligonucleotide primers. Sequences of primers used in this study for amplification of target sequences. Primer names, target sequences and nucleotide sequences (5′–3′) are provided. All primers were synthesized commercially.

**Table S3:** integration of specific quantifier and qualifier ions for lignin subunits. Qualifier and quantifier ions selected for lignin subunit (H, G, and S) identification and quantification in GC–MS analyses. Listed are the monitored ions (m/z) and their retention time (RT). RF= response factor

| Compound | Quantifier<br>(*m/z*) | Qualifier 1<br>(*m/z*) | Qualifier 2<br>(*m/z*) | Qualifier 3<br>(*m/z*) | mean RT<br>(min) | RF |
| --- | --- | --- | --- | --- | --- | --- |
| Tetracosane | 71.0855 | 85.1011 | 97.1011 | 113.1324 | 22.5808 | - |
| H (erythro) | 239.0917 | 189.0726 | 205.1035 | - | 23.9424 | 0.1950 |
| H (threo) | 239.0918 | 189.0726 | 205.1035 | - | 24.0379 |  |
| G (erythro) | 235.1143 | 269.1025 | 295.1179 | 403.1248 | 25.7936 | 2.4803 |
| G (threo) | 235.1145 | 269.1027 | 295.1180 | 403.1247 | 25.9128 |  |
| S (erythro) | 265.1255 | 299.1135 | 325.1289 | 433.1356 | 27.8152 | 1.9849 |
| S (threo) | 265.1254 | 299.1136 | 325.1287 | 433.1359 | 27.9878 |  |

**Table S4:** MRM transitions for cis- and trans-cinnamic acid. Qualifier and quantifier ions selected for *cis*- and *trans*-cinnamic acid identification and quantification in GC–MS analyses. Listed are the monitored ions and their use as quantifier or qualifier ions for confirmation of compound identity and analytical reliability. CE = collision energy

| Compound | RT<br>(min) | Quantifier/Qualifier ion | Precursor ion | Product ion | CE |
| --- | --- | --- | --- | --- | --- |
| *cis*-cinnamic acid | 11.82 | Quantifier | 205 | 131 | 20 |
|  |  | Qualifier | 205 | 161 | 5 |
|  |  | Qualifier | 205 | 103 | 20 |
|  |  | Qualifier | 161 | 59 | 20 |
| *trans*-cinnamic acid | 13.23 | Quantifier | 205 | 131 | 20 |
|  |  | Qualifier | 205 | 161 | 5 |
|  |  | Qualifier | 205 | 103 | 20 |
|  |  | Qualifier | 161 | 59 | 20 |

